# Mimicking an *in cellulo* environment for enzyme-free paper-based nucleic acid tests at the point of care

**DOI:** 10.1101/2024.02.27.582375

**Authors:** Jeffrey W. Beard, Samuel L. Hunt, Alexander Evans, Coleman Goenner, Benjamin L. Miller

## Abstract

Point of care (PoC) nucleic acid amplification tests (NAATs) are a cornerstone of public health, providing the earliest and most accurate diagnostic method for many communicable diseases, such as HIV, in the same location the patient receives treatment. Communicable diseases disproportionately impact low-resource communities where NAATs are often unobtainable due to the resource intensive enzymes that drive the tests. Enzyme-free nucleic acid detection methods, such as hybridization chain reaction (HCR), use DNA secondary structures for self-driven amplification schemes producing large DNA nanostructures and capable of single molecule detection *in cellulo*. These thermodynamically driven DNA-based tests have struggled to penetrate the PoC diagnostic field due to their inadequate limits of detection or complex workflows. Here we present a proof-of-concept NAAT that combines HCR-based amplification of a target nucleic acid sequence with paper-based nucleic acid filtration and enrichment capable of detecting sub pM levels of synthetic DNA. We reconstruct the favorable hybridization conditions of an *in cellulo* reaction *in vitro* by incubating HCR in an evaporating, microvolume environment containing poly(ethylene glycol) as a crowding agent. We demonstrate that the kinetics and thermodynamics of DNA-DNA and DNA-RNA hybridization is enhanced by the dynamic evaporating environment and inclusion of crowding agents, bringing HCR closer to meeting PoC NAAT needs.

## INTRODUCTION

Long existing healthcare disparities in resource-limited regions have been exacerbated over the years by globalization^1^ and climate change,^2^ increasing the need for point-of-care (PoC) tests to reach remote regions and prevent wide-spread infections in vulnerable populations.^3^ For example, human immunodeficiency virus (HIV) is a global concern disproportionately affecting residents of low-resource communities.^4^ Individuals newly infected with HIV (acute phase) have high viremia and need to start antiretroviral therapy as soon as possible to improve their health outcome and reduce transmissibility.^5^ Because acute HIV is pre-seroconversion, or prior to the presence of detectable levels of anti-HIV antibodies, it can only be accurately detected using a nucleic acid amplification test (NAAT).^6^ Unfortunately, there are no PoC NAATs for detecting acute phase HIV, which means infected individuals can go weeks to months without knowing they are infected.

Enzymatic methods for nucleic acid amplification have revolutionized modern diagnostics (as well as modern biology), but the cost and environmental sensitivity of enzymes can cause problems with their use in resource-limited areas. Enzyme-based NAATs like reverse transcription polymerase chain reaction (RT-PCR)^7^ and loop-mediated isothermal amplification (LAMP)^8^ are the most widely used methods for detecting HIV RNA or proviral DNA for acute HIV diagnosis. While these NAATs are highly sensitive, they are also largely inaccessible in low-resource settings due to their reliance on clinical environments and trained operators.^9,10^ Automated RT-PCR instruments such as the GeneXpert by Cepheid Inc. and the m-PIMA by Abbott Inc. are resource-demanding and require significant training to operate, making them challenging to use at PoC in low-resource settings.^11^ LAMP has become a popular alternative to RT-PCR because its amplification enzymes are more robust and operate isothermally. However, LAMP’s use of multiple sets of primers has been reported to result in unacceptable rates of off-target amplification outside of clinical settings.^12–14^ CRISPR is another enzyme-driven detection method that has been the subject of considerable study for PoC nucleic acid detection, but it must be coupled with a NAAT such as recombinase polymerase amplification (RPA) to be effective.^15–17^ CRISPR-RPA-based NAATs have been successful for PoC saliva tests but have struggled to penetrate the PoC sector for blood tests, as is needed for acute HIV diagnosis, because of the enzymes’ propensity for off-target amplification^18^ or inactivation in a non-sterile environment.^19^

Alternatives to enzyme-based amplification that instead rely on oligonucleotide self-assembly and self-processing are therefore of significant interest. A particular example that would seem ideally suited for diagnostic use is hybridization chain reaction (HCR). HCR takes advantage of the programmability of DNA molecules to harness potential energy in metastable hairpin structures that can fuel the assembly of massive double stranded DNA nanostructures when in the presence of a trigger sequence.

HCR was first reported in 2004 by Dirks and Pierce^20^ and involves a minimum of two ssDNA sequences engineered to form individual metastable hairpins, each with strategically located complementary regions that will promote a cascade of hairpin-to-hairpin hybridization when in the presence of a target nucleic acid sequence (Figure 1a). In controlled laboratory settings, fluorophore-conjugated HCR probes have been used for mRNA detection *in cellulo* with single-copy sensitivity.^21^ *In vitro*, HCR coupled with electrochemical sensors have demonstrated higher (but still impressive) limits of detection for DNA and miRNA targets with single aM analytical sensitivity,^22–24^ but the workflows for these sensors require complicated multi-step procedures and sterile operating conditions unsuitable for PoC applications. The *in cellulo* example and *in vitro* electro-chemical sensor examples can reach these extraordinarily low limits of detection in part by using multiple wash steps to remove unused HCR DNA hairpins to reduce background noise, a procedure that adds unwanted complexity to a PoC test.

**Figure 1.**
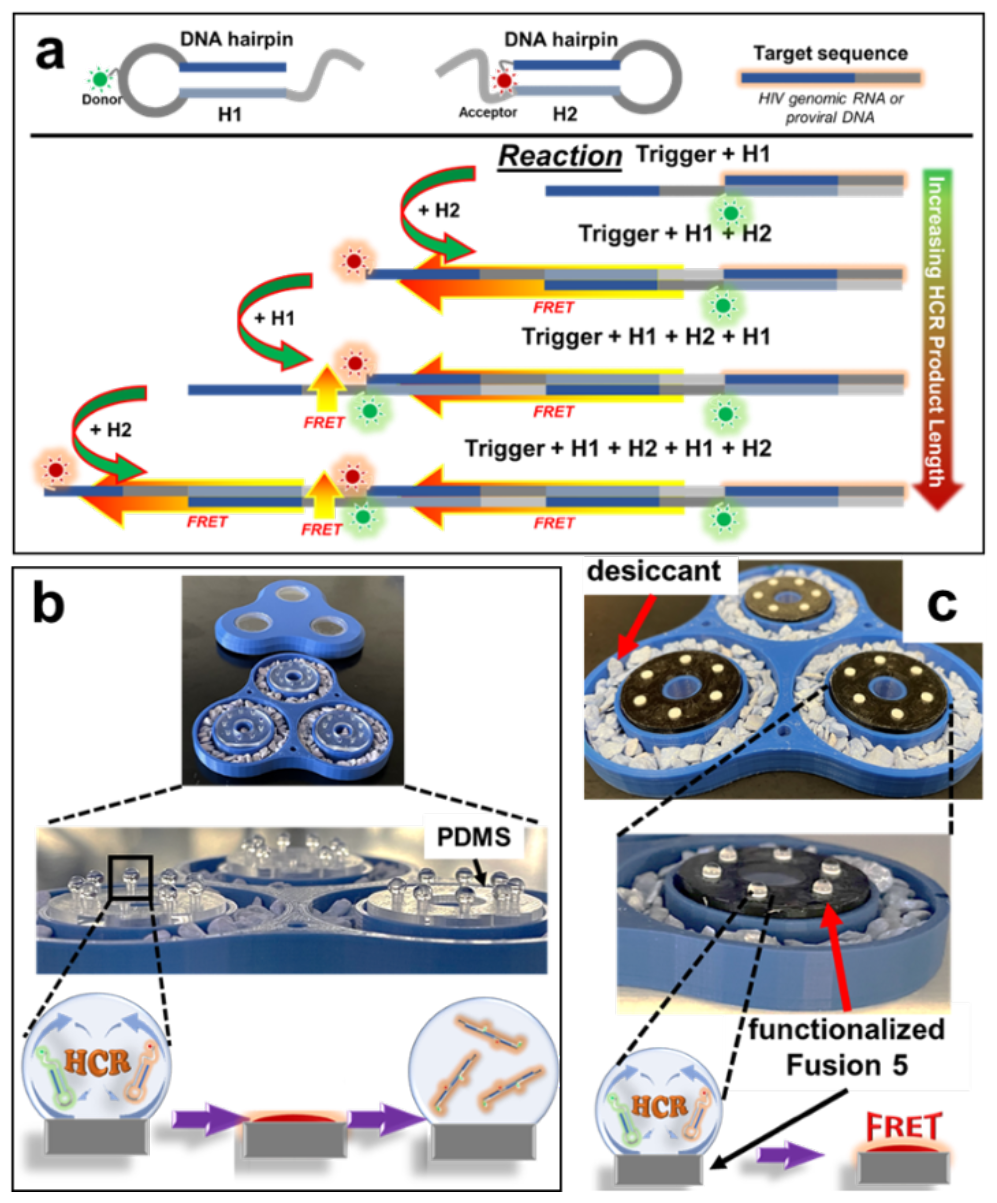
HCR scheme and sessile droplet experimental setups. a) Schematic of HCR using FRET (Cy3 acceptor and Cy5 donor fluorophores) for fluorescence detection of a target initiator sequence. H1 has a complementary region to the initiator sequence, causing it to unfold from its secondary structure and hybridize with the initiator, exposing a complementary sequence to H2. H2 hybridizes with H1, exposing a complementary sequence to H1, continuing the cascade. b) and c) HCR incubated in a sessile droplet on PDMS pedestals (b) and nucleic acid capture filter (c) and evaporated in an enclosed drying chamber (blue) surrounded by desiccant. Evaporated droplets are rehydrated in (b) or left evaporated (c) for analysis.

HCR with fluorescence readout and without a wash step has proven less sensitive, yielding tens of pM detection limits when carried out at an elevated temperature of 37 °C with synthetic DNA target.^25,26^ Combining HCR with enzymatic amplifications such as LAMP^27^ and CRISPR^28^ has enabled detection limits of 30 RNA copies and 1.5 fM, respectively. These examples demonstrate the versatility of HCR and its ability to be adapted towards *in vitro* detection of nucleic acids. The ideal PoC HIV test would be simple in its functional form but have detection limits nearing single molecule detection.

We set out to narrow the gap between the detection limit required for a PoC HIV test and what has been demonstrated thus far for HCR *in vitro*. Because HCR can detect a single copy of an mRNA inside a cell, we hypothesized that molecular crowding might contribute to the efficiency of *in cellulo* HCR and the extraordinarily low detection limit. To mimic the cell’s crowded environment,^29^ we added poly(ethylene glycol) (PEG) to the HCR hybridization buffer. PEG has been shown as an effective crowding agent to modify and stabilize conformational changes in DNA hairpins.^30^ We also allowed the reaction to evaporate in a sessile droplet (Figures 1b and 1c), which takes advantage of concentration enhancement as the solution dries and results in a nearly two-dimensional plane that can be easily viewed using a fluorescence microscope. To demonstrate HCR’s potential for low-resource PoC nucleic acid testing, we combined this approach with capture of synthetic HIV RNA and synthetic proviral DNA on chitosan-functionalized filter paper, as this material has been demonstrated to electrostatically adsorb and concentrate nucleic acids from a sample with high efficiency.^31–33^ By coupling this low-cost and user-friendly nucleic acid extraction method with fluorescently tagged HCR probes, we report what is to our knowledge the first example of enzyme-free nucleic acid detection directly on filter paper.

## RESULTS AND DISCUSSION

### DESIGNING HCR HAIRPINS AGAINST AN HIV GENOMIC TRIGGER

We designed HCR hairpins to target HIV using guidelines described by Ang and Yung.^26^ Their best-performing HCR hairpins had a stem length of 12 nucleotides, 66% guanine and cytosine (GC) content, and a 6-nucleotide sticky toehold region with 33% GC content. With these parameters in mind, we identified a GC rich (50% GC content) 18-nucleotide sequence, hereafter referred to as the “trigger”, from the conserved Gag gene region of HIV-1 vector pNL4-3. Our resulting hairpin designs designated H1 and H2 consisted of a 13-nucleotide stem with 53.8% GC content and a 6-nucleotide toehold region with a 33% GC content (Table S1).

Initial experiments confirmed that combining trigger, H1, and H2 yielded HCR products. Trigger concentration was found to have a notable impact on the molecular weight of polymers produced from H1-H2 assembly. At higher trigger concentrations (lanes 2 and 3 of Figure 2a) the molecular weights of the resulting HCR products are generally smaller than what is observed at lower trigger concentrations (lanes 3 and 4 of Figure 2a). The stunted HCR product in the presence of high trigger concentrations is the result of rapid H1 depletion due to H1-trigger hybridizations, producing short-length HCR products.^34^ This phenomenon is dependent on the ratio of hairpins to trigger availability and is consistent across a variety of hairpin and trigger concentrations (Figure S5).

**Figure 2.**
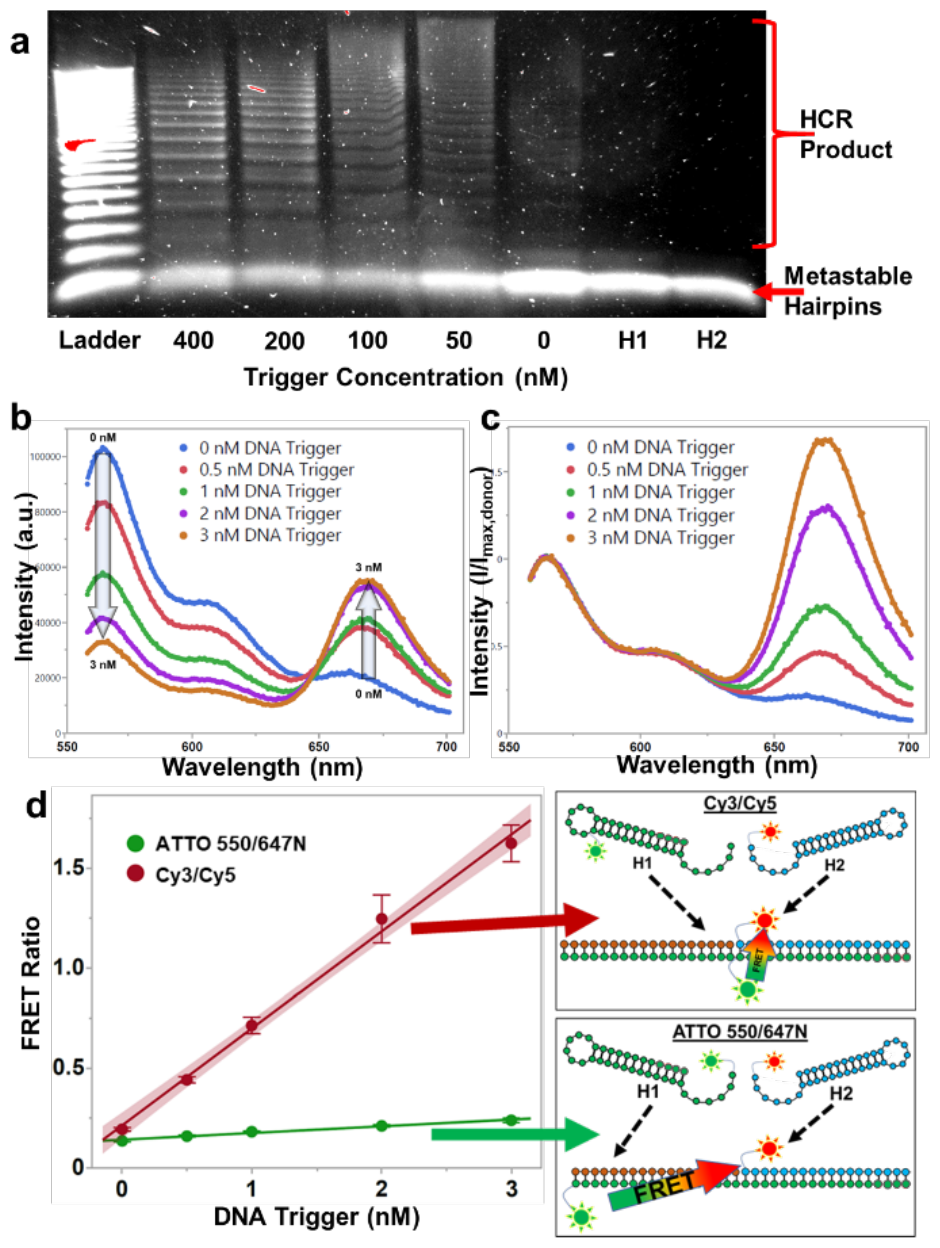
Validation of FRET-HCR hairpins against an HIV nucleic acid target. a) Agarose gel showing 500 nM concentrations of HCR hairpins incubated with synthetic proviral DNA trigger at different concentrations and a 20-base pair DNA ladder. b) Spectra of reactions using 5 nM H1-Cy3 and H2-Cy5 hairpins and varying synthetic DNA trigger concentrations. c) Normalized spectra from (b). d)FRET ratio of ATTO 550/647N dyes and Cy3/Cy5 dyes HCR scheme against synthetic DNA trigger concentrations with 5 nM hairpin concentrations. To the right are cartoon illustrations of the two FRET schemes. All panels used 5X SSC buffer with 0.05% Tween 20 and were incubated in microcentrifuge tubes for 1 hour. Shaded regions represent the 95% confidence intervals.

To quantify the presence of a trigger sequence, the emission spectrum of the reaction was measured (Figure 2b) and normalized to the peak donor intensity (Figure 2c). The normalized H2 peak (FRET ratio), was used to quantitatively assess buffer optimization and reaction conditions. We compared two pairs of donor and acceptor fluorophores: ATTO 550 with ATTO 647N, and Cy3 with Cy5. ATTO dyes are reportedly more stable and have higher quantum output than commonly used carbocyanines;^35^ however, the linking chemistry used by the commercial supplier used here limits the ATTO fluorophore conjugations to the 5’ and 3’ termini of the DNA hairpin oligos. This is a disadvantage in our HCR-FRET scheme, resulting in a loss of about half of the FRET energy transfer due to a distance of 18 nucleotides between donor and acceptor fluorophores in the HCR product (Figure 2d). The internal placement of the Cy3 fluorophore in H1 resulted in a FRET ratio 292% greater than the ATTO dyes at 1 nM DNA trigger concentration, and as such, we chose to use the Cy3/Cy5 scheme for the majority of our experiments.

### SESSILE DROPLET INCUBATION

With an initial demonstration of a working HCR for HIV DNA in hand, we next set out to identify methods for enhancing the detection limit. To recapitulate the optimal hybridization conditions found *in cellulo*, we tested two possibilities: sessile droplet incubation, and inclusion of crowding agents, both separately and together. Sessile droplets have been shown to provide a dynamic small volume environment for enhancing enzymatic activity^36^ We hypothesized that sessile droplet reactions could likewise enhance HCR detection limits by mimicking the enclosed, dynamic cellular environment that enables single-copy nucleic acid sensitivity for HCR. While microdroplet incubations immersed in oil are known to improve the efficiency of NAATs,^37–39^ and Yen et al. showed a high efficiency of DNA duplex hybridization in oil encapsulated pL volume droplets,^40^ these studies do not leverage the advantages of an evaporating system. As the liquid evaporates, the volume of the sessile droplet decreases, increasing the concentrations of all components in the system and changing the hybridization interactions.

Evaporation of a sessile droplet increases the concentration of trigger over time, thus favoring initiation of the HCR. One potential concern, however, was that time-dependent increases in the concentration of other species could have a negative effect on the process. For example, 5X SSC is a commonly used hybridization buffer in HCR experiments^26,41–43^ for its high sodium content (750 mM sodium chloride) which is favorable for nucleic acid hybridization. However, in an evaporating environment the increasing salt concentration impacts the metastability of the DNA hairpins (Figure 3a). As water evaporates from the 5 μL starting sample of 0.5X SSC buffer (75 mM sodium concentration) the buffer composition increases in concentration. The buffer eventually concentrates to 5X SSC (750 mM sodium concentration) once the volume reaches 0.5 μL, shifting the measured melting temperature of H1 from 73 °C to 84 °C and demonstrating the importance of a diluted starting buffer. A starting sodium concentration greater than 75 mM was observed to over-stabilize the DNA hairpins early in the evaporative incubation and result in a retardation of hybridization events (Figure 3b, purple). Another important component of the evaporating droplet buffer is MgCl_2_. The addition of MgCl_2_ to the buffer reduced the background noise from uninitiated HCR products (Figure 3b, green and red); this is likely due to structural stabilizing effects of the divalent cations on the metastable hairpins.^44^

**Figure 3.**
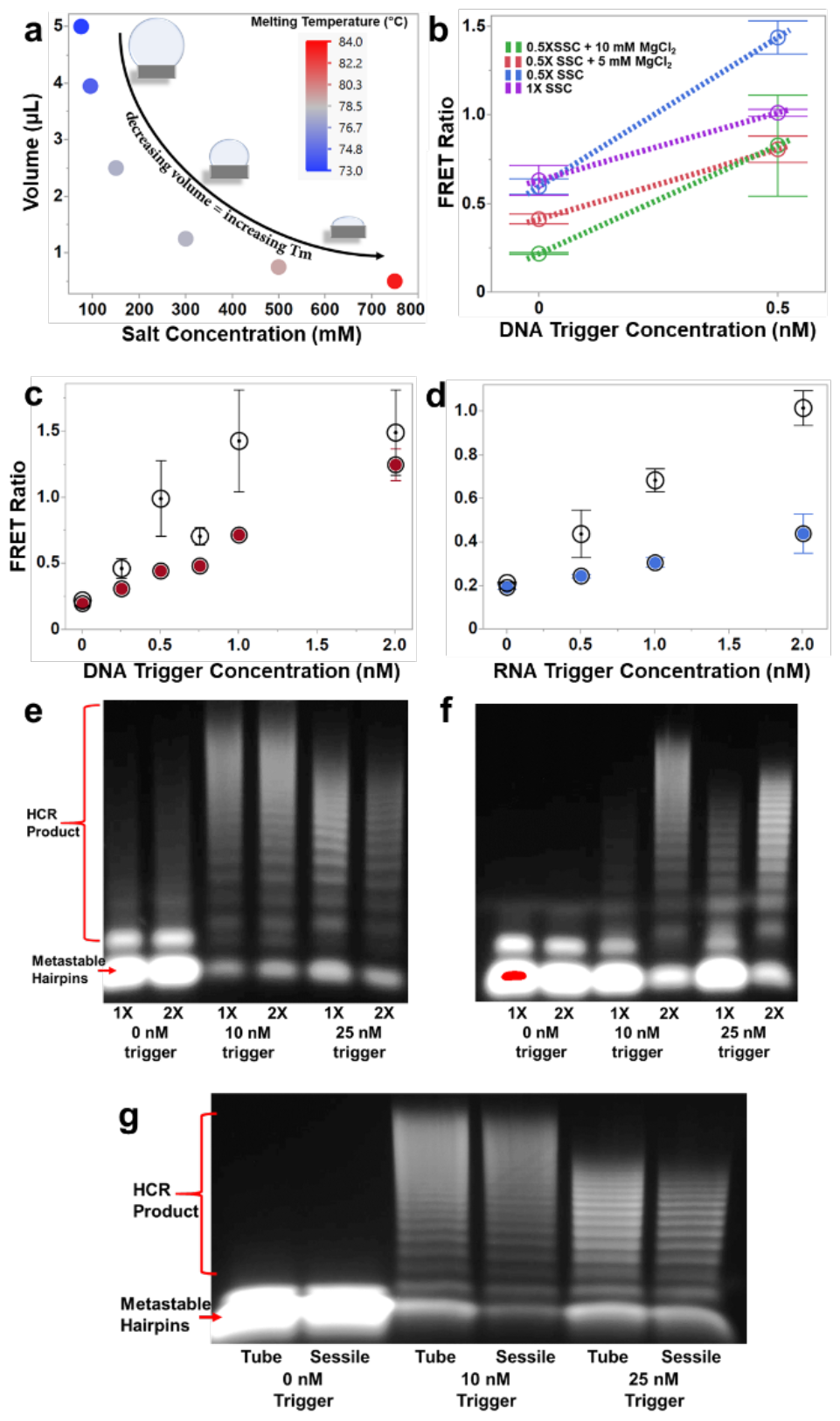
Incubating HCR in sessile droplets. a) Experimental data of H1 melting temperatures (colorimetric scale) at different sodium concentrations found in 0.5X SSC to 5X SSC (x-axis) and plotted against evaporating volume of a theoretical sessile droplet (y-axis). b) Cy3/Cy5 FRET ratios with 0 nM or 0.5 nM synthetic DNA trigger incubated for 1 hour in sessile droplets with different buffer conditions. Each buffer includes 0.1% Tween 20. c) and d) Cy3/Cy5 FRET ratio versus synthetic DNA trigger (c) and synthetic RNA trigger (d). 5 nM hairpins were incubated for 1 hour in sessile droplets (open circles) in 0.5X SSC and 10 mM MgCl_2_ buffer with 0.1% Tween 20 (c) or 0.05% Tween 20 (d), and micro-centrifuge tubes (colored circles) incubated in 5X SSC with 0.05% Tween 20. In (d), RNA microcentrifuge tube incubations (blue circles) were incubated for 90 minutes. e) Agarose gel of sessile droplet incubations using 50 nM Cy3/Cy5 hairpins. 1X lanes consist of 5 μL sessile droplet incubations for 60 minutes in 0.5X SSC with 10 mM MgCl_2_ and 0.05% Tween 20. The 2X lanes are two times the concentration of everything in 1X including trigger concentration and buffer but incubated in 2.5 μL droplets for 45 minutes. 2X lanes had double the DNA trigger concentration labeled below each pair of lanes (e.g. the 2X lane contained 20 nM trigger but is being compared to the 10 nM trigger of the 1X lane). f) The same experiment as (e) but incubated in cuvettes instead of sessile droplets. g) Agarose gel of HCR product from cuvette and sessile droplet incubations in 5X SSC with 0.05% Tween 20 buffer. The volume of the sessile droplet was maintained at approximately 5 μL for the entirety of the hour by adding 1.25 μL of Nanopure water to the evaporating droplet every 10 minutes. Gels were imaged using the Cy5 channel. Error bars show standard deviation from an n = 3, and shaded regions represent the 95% confidence intervals.

We observed a 3-fold improvement in limit of detection (LOD) for DNA-triggered HCR by the sessile droplet incubation, and the LOD for the RNA-triggered reaction was improved by over 10-fold (Figures 3c and 3d and Table S3). We tested the specificity of the HCR-sessile droplets by incubating the reactions with excess salmon sperm DNA (Figure S7) and yeast tRNA (Figure S8). We found the HCR reaction was not hindered or exacerbated by off-target hybridization events, indicating our HCR-sessile droplet scheme is highly specific to its target sequence.

Reactions in the sessile droplet using synthetic DNA trigger resulted in significantly more HCR product than reactions using equivalent concentrations of synthetic RNA trigger (Figures 3c and 3d). It has been shown that DNA:RNA hybrids have lower thermodynamic stability than DNA:DNA duplexes,^45^ which may thermodynamically impair the ability of H1 to unfold from its secondary hairpin structure and initiate a cascade. However, the RNA trigger required a 90-minute incubation while the DNA trigger was only incubated for 60 minutes, indicating the reduction in HCR product using RNA trigger is likely a kinetic issue. Reduced kinetics between DNA:RNA hybridization compared to DNA:DNA hybridization have been observed by other groups.^46^

To understand the driving force behind the improved HCR in sessile droplets, we tested how increasing the concentrations of the hairpins, trigger, and buffer components during evaporation impact the reaction. Two sets of sessile droplet reactions were run: a “1X” reaction in a 5 μL droplet, and a “2X” reaction in a 2.5 μL droplet, the latter mimicking the reagent concentrations and volume of the 1X droplet after 15 minutes of evaporation. Gel electrophoresis at the conclusion of both reactions revealed nearly identical product formation (Figure 3e), indicating most hybridization events take place sometime after half of the initial volume has evaporated.

As the sessile droplet evaporates, the trigger concentration increases at the same rate as the hairpins, eventually reaching optimal hairpin and trigger concentrations for maximum product output with minimal background noise. We have found that increasing the concentration of the DNA hairpins without increasing the trigger concentration produces comparable HCR product but results in an excess of un-used, metastable hairpins that contribute to background noise (Figure S6). When the same 1X and 2X reactions as described above were incubated in non-evaporating conditions (Figure 3f), the 2X reaction produced significantly more HCR product than the 1X reaction, showing that concentrating the hairpins and trigger at the same rate is an important aspect to the success of the HCR-sessile droplet incubation.

An additional potential driving force to consider behind the improved HCR kinetics within the sessile droplet microenvironment is Marangoni flow. This is an interfacial phenomenon produced by a surface tension gradient at the interface of two fluids and has been demonstrated in sessile droplets using gradients of volatile vapors,^47^ temperature,^48^ and detergents^49^ to produce transverse “mixing” vortices within the evaporating droplet. These vortices have been demonstrated to overcome traditional capillary flow, which is a lateral, non-mixing flow responsible for the well-known coffee ring effect in evaporating droplets.^50^

We tested the influence of Marangoni flow in sessile droplet HCR by comparing sessile droplet HCR to a cuvette incubation using 5X SSC with 0.05% Tween 20 buffer for both scenarios, and identical starting concentrations of trigger and hairpins. The sessile droplet volume was replenished to its original 5 μL volume every 10 minutes with Nanopure water for 1 hour to prevent positive effects from concentrating the reagents. If Marangoni flow was influencing the hairpin hybridization kinetics, we would expect to see increased HCR product in the sessile droplet over the microcentrifuge tube incubation even though the sessile droplet was never allowed to decrease significantly in volume. But this was not observed (Figure 3g): nearly identical HCR product was obtained in both the sessile and microcentrifuge tube incubations. As such, it does not appear that Marangoni flow is an influencing factor in sessile droplet HCR and further supports the theory that the observed enhancement is primarily due to the increasing reagent concentrations during evaporation.

### CROWDING AGENTS IN SESSILE DROPLET INCUBATIONS

During initial buffer optimization experiments, we observed a significant impact of Tween 20 on HCR product formation (Figure 4a), which is in part due to an increase in fluorescence output from the H1 fluorophore (Figures 4b and S5). A possible explanation for the increase in fluorescence output is reduction of guanine quenching. Guanine is reported to have strong quenching effects on ATTO dyes ^51^ and mild quenching effects on carbocyanine dyes,^52^ and the amphiphilic Tween 20 may provide a protective barrier around the slightly hydrophobic fluorophores.

**Figure 4.**
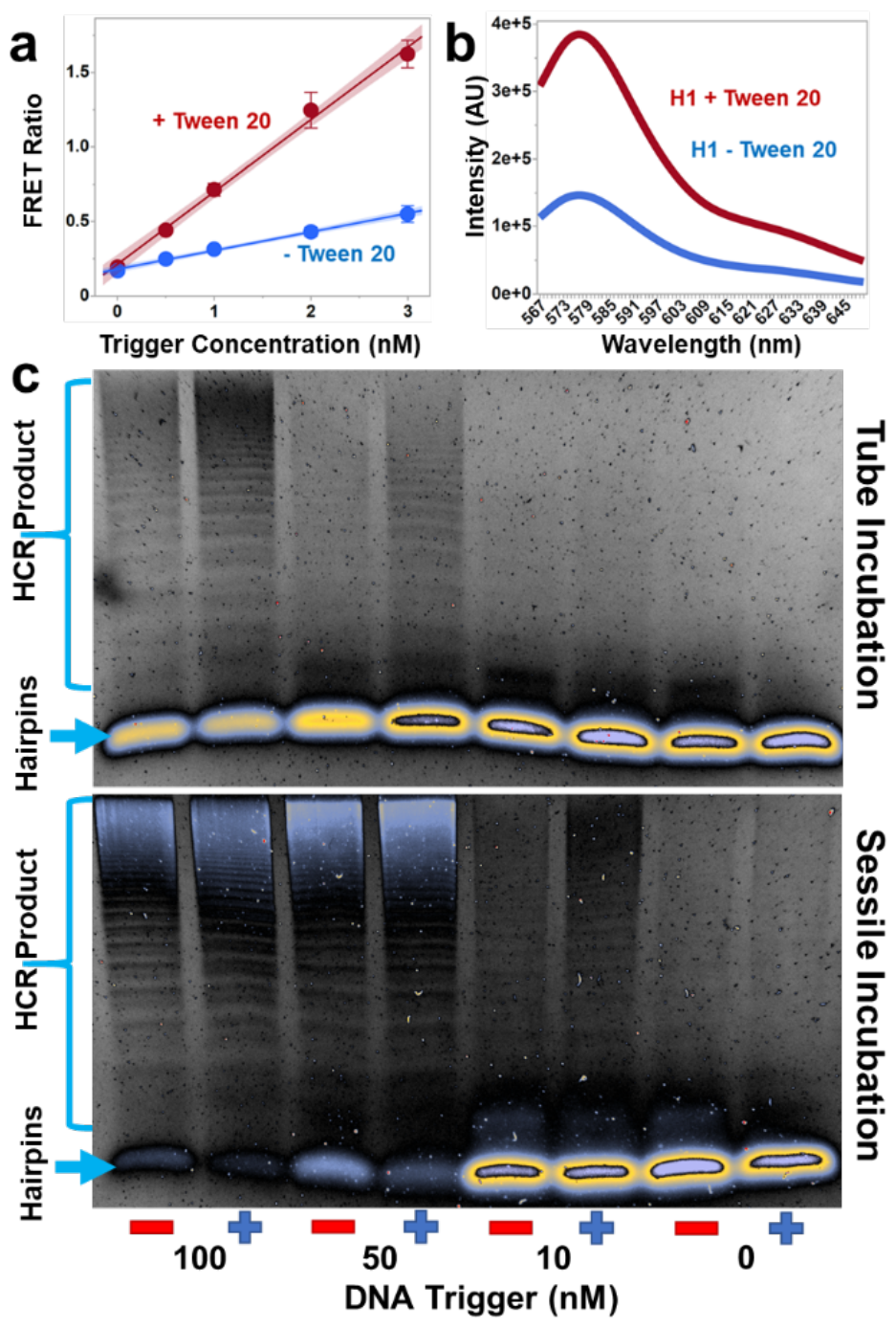
Tween 20’s impact on HCR. a) Cy3/Cy5 FRET ratios of hairpins incubated with synthetic DNA trigger for 1 hour in micro-centrifuge tubes with (red) and without (blue) the presence of 0.05% Tween 20. b) ATTO 550 spectra with (red) and without (blue) the presence of 0.05% Tween 20. c) Agarose gel stained with SybrSafe showing 500 nM hairpins (no fluorophores) incubated with synthetic DNA trigger in microcentrifuge tubes (top) and sessile droplets (bottom) with (+) and without (-) the addition of Tween 20 in their respective hybridization buffers. Error bars show standard deviation from an n = 3, shaded regions represent the 95% confidence interval.

However, enhanced fluorescence output is not the only explanation for the observed increase in FRET intensity. We found that Tween 20 also improves the formation of HCR product in both the microcentrifuge tube and the sessile droplet incubations (Figure 4c). We hypothesized that the improved hybridization could be from a crowding effect imposed on the nucleic acids by the Tween 20. Tween 20 is largely composed of PEG, which has been shown to modify and stabilize conformational changes in DNA hairpins via a molecular crowding effect.^30^ We tested this hypothesis by adding PEG to the microcentrifuge tube reaction buffer (5X SSC) with and without Tween 20 (Figure 5a). Reactions containing a 10% w/v PEG (with and without Tween 20) produced significantly higher molecular weight product (i.e., longer HCR polymers) than reactions without PEG.

**Figure 5.**
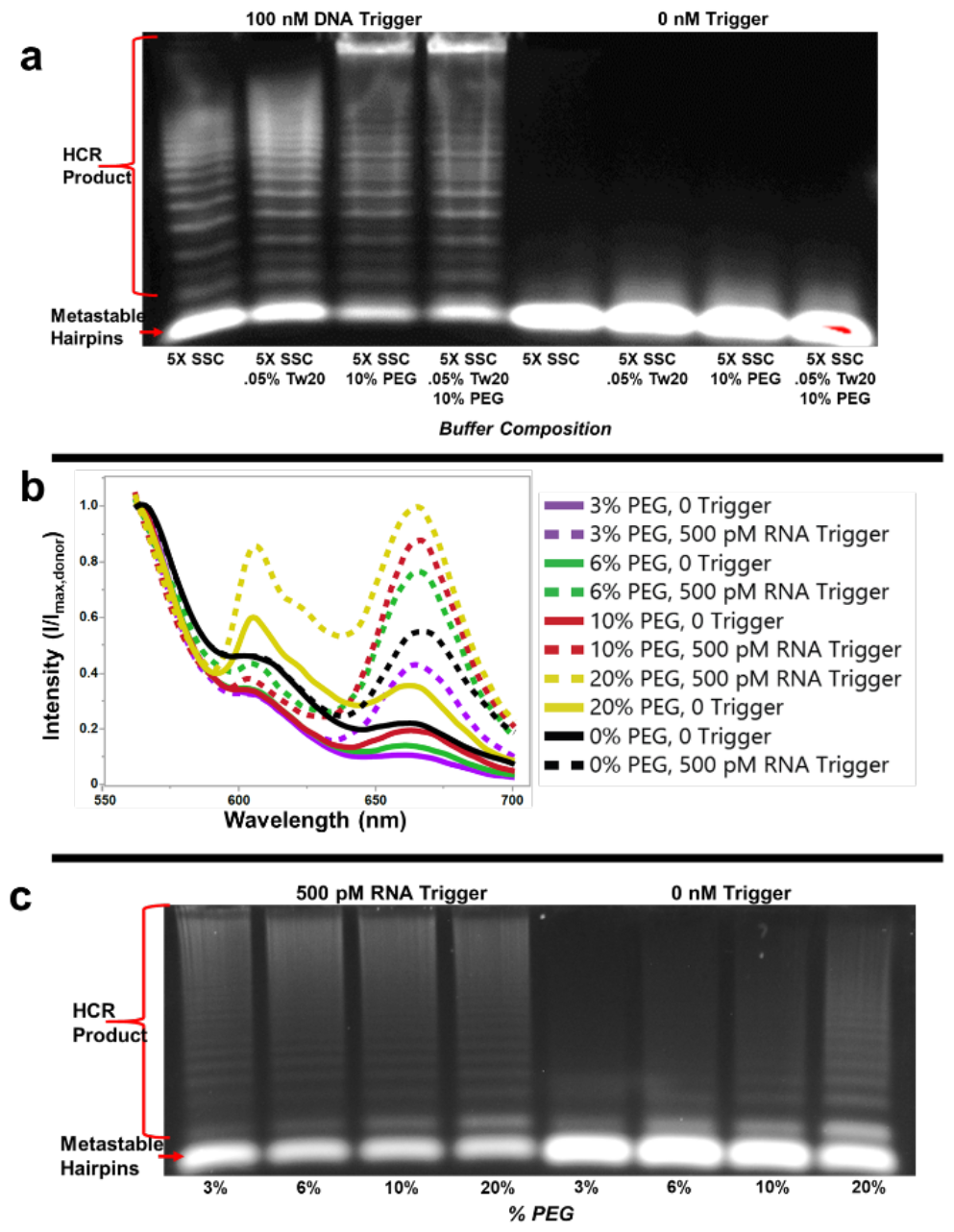
Crowding enhances HCR but also increases the uncatalyzed reaction at high PEG concentration. a) Agarose gel showing HCR product from 500 nM Cy3/Cy5 hairpins incubated in micro-centrifuge tubes with and without synthetic DNA target in different buffer compositions. b) Normalized Cy3/Cy5 HCR spectra of 5 nM hairpins incubated with and without synthetic RNA trigger in sessile droplets. Buffers were 0.5X SSC with 10 mM MgCl_2_ and 0.05% Tween 20 with varying amounts of PEG by % w/v. c) The equivalent experiment described in (b) but imaged in an agarose gel.

To further test the effect of PEG on the HCR-sessile droplet, we incubated HCR reactions containing RNA trigger with 0%, 3%, 6%, 10%, and 20% w/v PEG in evaporating sessile droplets and compared the resulting FRET spectra (Figure 5b). The difference in intensity between the 500 pM RNA trigger experiment and the no-trigger control experiment was consistently greater in the presence of PEG compared to the 0% w/v PEG experiment (Table 1). Comparing the difference in intensities between the trigger and no-trigger experiments in Table 1, the 6%, 10%, and 20% w/v PEG experiments performed the best with a 193%, 210%, and 201% improvement, respectively, over the 0% w/v PEG experiment. However, increasing the PEG content of the buffer also resulted in an increase in uninitiated hybridization events in the no-trigger controls (Figures 5b and 5c).

**Table 1.** Normalized FRET intensities from Figure 5b.

| % PEG | 0% | 3% | 6% | 10% | 20% |
| --- | --- | --- | --- | --- | --- |
| 500 pM RNA Trigger | 0.529 | 0.418 | 0.737 | 0.845 | 0.977 |
| 0 Trigger Control | 0.219 | 0.105 | 0.138 | 0.194 | 0.353 |
| Difference | 0.310 | 0.313 | 0.599 | 0.651 | 0.624 |

### ENZYME-FREE FILTRATION AND DETECTION OF NUCLEIC ACIDS

Capture of DNA on paper filters is well known. In particular, the use of chitosan-functionalized Fusion 5 filter has been demonstrated to electrostatically adsorb DNA with high efficiency.^31,33^ We anticipated that functionalized Fusion 5 could serve as the end point of a viral lysis and nucleic acid isolation strategy, but first needed to confirm that adsorbed nucleic acids would still efficiently initiate HCR. Before testing capture of nucleic acids on a filter, we first tested our HCR-sessile droplet reaction on the filter material. Synthetic DNA or RNA trigger sequences were applied to filters as 2 μL droplets and evaporated prior to adding the PEG-enhanced HCR sessile droplet mixture. FRET intensities of the final HCR products were quantified by fluorescence microscope. As expected, the DNA trigger yielded the greatest intensity increase over background with high statistical significance at 5 attomoles of trigger (Figure 6a). The RNA trigger experiments were less efficient, showing significant increase in intensity at 100 attomoles of target (Figure 6b and 6c). This may be from the less favorable conditions of DNA:RNA hybridization versus DNA:DNA hybridization, as previously described.

**Figure 6.**
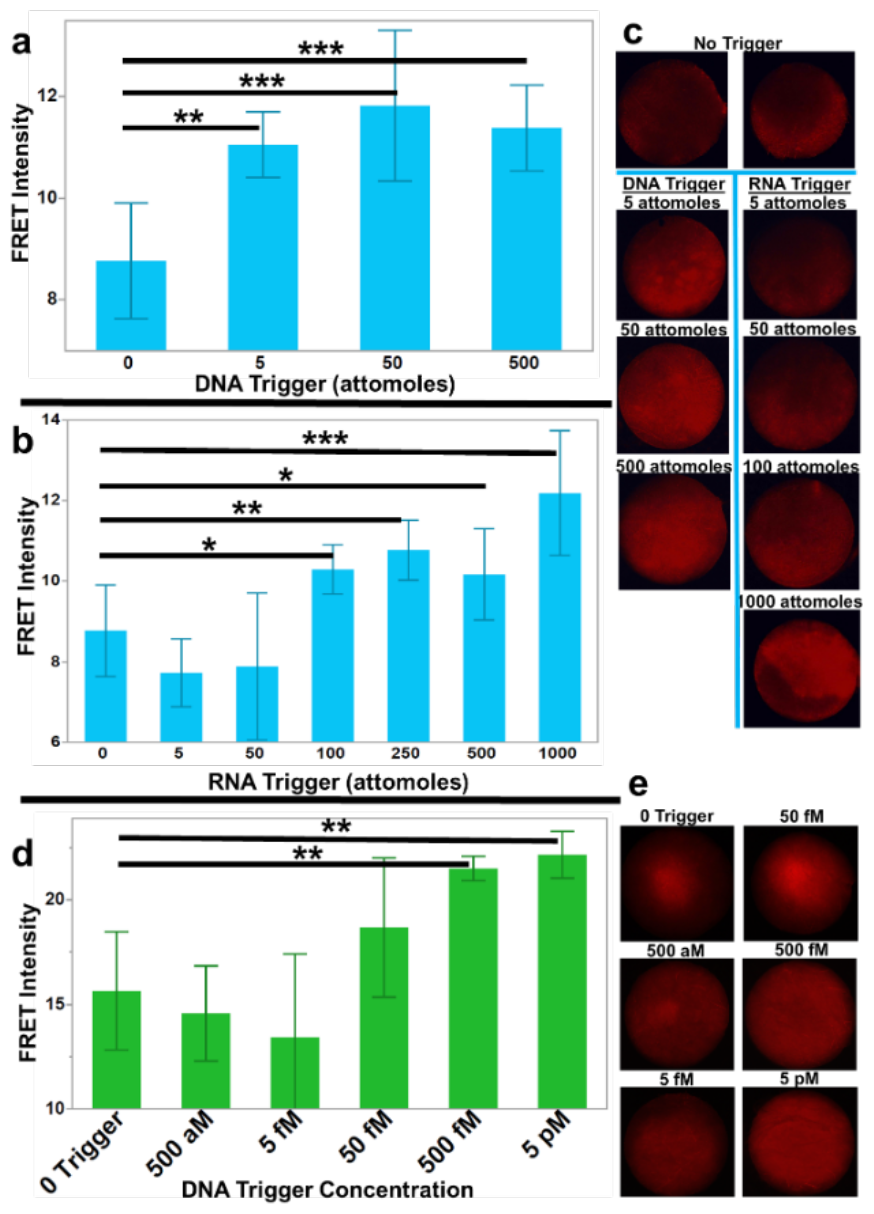
Fluorescence intensities of HCR product on functionalized Fusion 5 filter. a) and b) Binned fluorescent intensities of FRET-HCR product from dried trigger experiments on capture filter using synthetic proviral DNA (a) or synthetic HIV RNA (b) trigger (n = 10 for 0 trigger controls, n = 4 for 5, 50, and 1000 attomoles in (b), and n = 6 for all others). c) Fluorescence microscope images of functionalized Fusion 5 filters with dried HCR reaction from (a) and (b). d) Binned intensities from capture and detection of nucleic acids experiments using synthetic proviral DNA (n=4). e) Fluorescence microscope images of functionalized Fusion 5 filters with dried HCR reaction from (d). Error bars show standard deviation and p values were calculated using a two-tailed Student’s t-test (* p < 0.05, ** p < 0.01, *** p < 0.001).

Finally, we combined HCR-based nucleic acid detection with filtration and capture of the synthetic trigger. For high efficiency capture of nucleic acids, the chitosan must be exposed to a pH below its pKa of ∼6.4.^53^ We found that diluting the DNA in 1X PBS at pH 5 resulted in a capture efficiency approaching 100% at low concentrations of DNA (Figure S2). We were able to detect the filtered DNA trigger accurately down to a concentration of 500 fM (Figures 6d and 6e). This demonstrates that our enhanced HCR system can be used in a workflow compatible with paper-based nucleic acid filtration and enrichment.

## CONCLUSION

This proof-of-concept work shows the capture, enrichment, and detection of synthetic DNA targets directly on paper down to sub-pM limits of detection. The workflow is simple and robust. As such, this approach should facilitate development of simple, robust nucleic acid assays for PoC environments. We will continue to work on improving our enzyme-free hybridization scheme by exploring different crowding agents and concentrations in our evaporating buffer conditions. Our FRET-based detection scheme is inefficient due to inefficient energy transfer, so we will also improve our detection limits by implementing a single fluorophore detection scheme. Of particular importance, the results described here highlight the role of molecular crowding and concentration on HCR. We anticipate this observation will prove useful in many other applications of nucleic acid self-assembly or DNA nanotechnology.

While observed detection limits are significantly improved relative to other reported HCR-based assays, we anticipate that the overall sensitivity can be further improved. First, the current FRET based detection scheme is subject to significant background due to bleed through of the fluorophores. Use of a single fluorophore plus quencher scheme (by analogy to well-known molecular beacons) may avoid this issue. Second, the lower limit of detection for RNA in these experiments was roughly an order of magnitude higher than for DNA. Since a goal of this work is direct detection of HIV, it is particularly important to improve this. Our future work will test use of a DNA trigger that is activated by the presence of an RNA target, which have been used successfully in similar RNA detection assays.^54^

### EXPERIMENTAL SECTION

#### General Materials

Deionized water was further purified to 18 MΩ using a NANOpure II Water System (“Nanopure water”; Barnstead Thermolyne Corporation, Ramsey, MN). Poly-dimethylsiloxane (PDMS) was made from 184 Silicone Elastomer Base and 184 Silicone Elastomer Curing Agent (Dow Chemical Company, Midland, MI) in a 7:1 ratio and cured under vacuum at 50 °C. Sodium chloride, magnesium chloride hexahydrate, RNaseZap™, UltaPure agarose, SyberSafe DNA gel stain, and SyberGold DNA gel stain were purchased from Thermofisher Scientific (Waltham, MA). Sodium citrate was purchased from Mallinckrodt (Staines-upon-Thames, UK), and Tween 20 and 10x TBE buffer from Bio-Rad (Hercules, CA). Poly(ethylene glycol) (PEG) (MW 07435) was purchased from Sigma-Aldrich (Munich, Germany). All 3D printed structures were designed using SolidWorks 2020 Student Edition and printed using 1.75 mm PLA Filament (Hatchbox 3D, Pomona, CA) on an Original Prusa i3 MK3S+ 3-D Printer (Prusa Research, Prague, Czech Republic). All statistical data was analyzed using JMP Pro 16.

#### Nucleic Acid Oligomers

All oligomers were purchased from Integrated DNA Technologies (Coralville, IA) as desalted lyophilisate, resuspended in TE buffer, and stored at 4 °C until use. Fluorescently conjugated hairpins and synthetic HIV RNA were HPLC purified by the commercial company. H1 and H2 hairpin secondary structures and thermodynamic stabilities were computationally verified using RNAstructure.^55^ DNA hairpins were annealed prior to each experiment using an Amplitron II thermocycler (Barnstead Thermolyne Corporation, Ramsey, MN) by heating to 95 °C for 5 minutes, then cooling to room temperature over 30 minutes. Hairpin melting temperatures were determined using 260 nm absorbance measured from 35-95 °C in 2 °C intervals in varying salt concentrations with a UV-1800 spectrophotometer (SHIMADZU, Kyoto, Japan). Hairpin secondary structures were experimentally verified by showing equivalent melting temperatures of hairpins in 5X SSC at 0.5, 1, and 2 μM concentrations (Figure S1). All Förster Resonance Energy Transfer (FRET) data was gathered using a Cy3 or ATTO 550 donor fluorophore conjugated to the H1 hairpin and a Cy5 or ATTO 647N acceptor fluorophore conjugated to the H2 hairpin (Table S1).

#### Agarose Gel and Spectrofluorometer HCR Experiments

The HCR reaction mixture consisted of a combination of H1 hairpins, H2 hairpins, and an initiator sequence of synthetic DNA or RNA (Figure 1a). Hairpin concentrations were 5 nM, 100 nM, or 500 nM depending on the experiment. HCR mixtures were incubated in 5 μL droplets on 1.5 mm diameter PDMS pedestals, enclosed in a 3D-printed drying device (Figure 1b, blue) surrounded by 3.2 grams of desiccant, or incubated in an enclosed microcentrifuge tube. The PDMS pedestals were designed to allow the droplets to form spherical shapes to maximize the area of the liquid-gas interface, and situated within the custom drying device to ensure consistent exposure to desiccant and drying conditions for each droplet. Glass slide covers were embedded in the top of the chamber to allow observation of pedestals without mechanical disturbance. Unless otherwise stated, all sessile droplet incubations were done using “sessile buffer” consisting of 0.5X SSC (75 mM sodium chloride and 15 mM sodium citrate), 10 mM MgCl_2_, and 0.1% or 0.05% Tween 20 for DNA and RNA trigger reactions, respectively. All buffers were pH 6.9 - 7.0 unless otherwise stated. Droplets were allowed to dry completely (∼60 minutes) in the custom drying device.

After drying, HCR products were rehydrated on their pedestals using 5 μL of Nanopure water and transferred to micro-centrifuge tubes or crystal cuvette for analysis via agarose gel electrophoresis or the spectrofluorometer, respectively. Rehydrated HCR products were diluted 3:1 in a 12% sucrose solution before being added to a 3% agarose gel (made with 1X TBE) and run in 0.5X TBE running buffer for 45 minutes at 130 volts. Gels were imaged using a ChemiDoc MP Imaging System (Bio-Rad Laboratories, Hercules, CA).

Spectrofluorometer analysis was performed using a Fluoromax 4 (Horiba, Kyoto, Japan) and consumed 30 μL of HCR product per cuvette. This was diluted in 70 μL Nanopure to fulfil the minimum volume requirements of the instrument and still have a detectable signal. Each FRET scheme was excited at the donor’s measured λ(max) and the emission spectral intensities captured. Cy3/Cy5 FRET schemes were excited at 550 nm and emission intensities from 558 to 700 nm were measured. FRET ratios were calculated by dividing the peak average acceptor fluorescence intensity (averaged from 664 to 669 nm) by the averaged peak donor fluorescence intensity (averaged from 562 to 567 nm). ATTO 550/647N FRET schemes were excited at a wavelength of 560 nm, and emission intensities from 568 to 700 nm were measured. FRET intensities were calculated by dividing the peak acceptor fluorescence intensity by the peak donor fluorescence intensity.

Non-sessile droplet reactions (microcentrifuge tube HCRs) were analyzed using the same volumes and dilutions as the sessile droplet experiments. All microcentrifuge tube HCRs were incubated for 1 hour at room temperature in 5X SSC with 0.05% Tween 20, unless otherwise stated.

#### Functionalizing and Testing Capture Filters

Whatman Fusion 5 (Cytiva, Marlborough, MA) was functionalized with chitosan (Sigma-Aldrich, Munich, Germany) using a protocol described by Rosenbohm et al.,^33^ and stored in a foil zipper lock bag at 4 °C until use. Capture efficiency of functionalized filters was tested using two random short DNA oligos (DNA oligo 1 and DNA oligo 2 in Figure S2d) or yeast tRNA (Sigma-Aldrich, Munich, Germany) by diluting the samples in 1X PBS (pH 5.0- 5.4, unless stated otherwise). Functionalized Fusion 5 was inserted into modified Slide-A-Lyzer™ MINI Dialysis Units (Thermofisher Scientific, Waltham, MA) and the diluted DNA or yeast tRNA was centrifuged through the filter units at 1300 x g or 4000 x g. Differences in the centrifugation speed did not impact capture results. Before and after filtration measurements of the eluate were taken using a NanoDrop™ Lite Spectropho-tometer (Thermofisher Scientific, Waltham, MA). A decrease in nucleic acid concentration in filtered eluate was used to indicated nucleic acid capture (Figure S2 and Table S2).

#### Detection of Nucleic Acids on Functionalized Filter

A 2 mm biopsy punch was used to punch 2 mm diameter disks of functionalized Fusion 5 filters. The filters were applied to double sided adhesive on the surface of 3D printed rings (Figure 1c, black rings) and placed in the drying chambers (Figure 1c, blue components). Synthetic DNA or RNA triggers were diluted in Nanopure water to respective concentrations and applied to the disks as 2 μL droplets where they evaporated over 15 minutes, leaving a finite number of target molecules.

The dried trigger on paper was resuspended in a 5 μL droplet of 5 nM H1 and H2 conjugated with Cy3 and Cy5, respectively, in 0.5X SSC with 10 mM MgCl_2_, 0.05% Tween 20, and 3% PEG. The reactions were fully evaporated for 60 minutes in the custom drying chambers described earlier and imaged at a gain of 4 and 100 ms exposure using a Cy3/Cy5 FRET cube and Olympus BX60 Microscope (Olympus Corporation, Tokyo, Japan) equipped with a SPOT Flex 64MP digital camera (Diagnostic Instruments Inc., Sterling Heights, MI). Images were analyzed using ImageJ 1.53c^56^ and results assessed by quantifying the binned intensity of each HCR paper disk.

#### Capture and Detection of Nucleic Acids on Functionalized Filters

Double sided adhesive (3M, Maplewood, MN) with 1.5 mm diameter holes was applied to the surface of a wicking layer (ShamWow®, Wallingford, CT). 2 mm disks of functionalized Fusion 5 filters were carefully adhered to the wicking layer, centered over the 1.5 mm holes in the adhesive. 1 mL of 1x PBS (pH 5) with varying concentrations of synthetic proviral DNA target was applied to the filters using a 1 mL syringe. The solution was wicked through the filters into the bottom layer (Figure S3).

The filters were then moved to double sided adhesive on the surface of custom 3D printed rings (Figure 1c, black rings) and dried for 10 minutes in the custom drying chambers (Figure 1b, blue component). H1 and H2 conjugated with Cy3 and Cy5, respectively, were diluted to 5 nM concentrations in 0.5X SSC with 10 mM MgCl_2_, 0.05% Tween 20, and 6% PEG (brought to pH 8 using Tris base) and were applied to the paper disks as 5 μL droplets. The PEG concentration was increased in these experiments to 6% due to further optimization of the crowding buffer concentrations. The reactions were fully evaporated for 60 minutes in the custom drying chambers and processed using the fluorescence microscope and FRET cube as described earlier with the exception that the images were taken at a gain of 4 and 500 ms exposure and images were converted to RGB color prior to analysis.

## Supporting information

Combined Supplementary Information

## SUPPORTING INFORMATION

Sequences and Tm experiments; functionalized Fusion 5 experiments; limit of detection calculations; HCR with varying hairpin concentrations; HCR with salmon sperm DNA and yeast tRNA; impact of Tween 20 on fluorescence. (PDF). The Supporting Information is available free of charge on the ACS Publications website.

## AUTHOR INFORMATION

**Corresponding Author**

^*^ **Benjamin L. Miller** *-Department of Dermatology, University of Rochester, Rochester, NY 14627, USA;*

**Authors**

**Jeffrey W. Beard** - *Department of Dermatology, University of Rochester, Rochester, NY 14627, USA*

**Samuel L. Hunt** - *Department of Dermatology, University of Rochester, Rochester, NY 14627, USA*

**Alexander Evans** - *Department of Biomedical Engineering, University of Rochester, Rochester, NY 14627, USA*

**Coleman Goenner** - *Department of Biochemistry and Biophysics, University of Rochester, Rochester, NY 14627, USA*

**Author Contributions**

The manuscript was written through contributions of all authors. All authors have given approval to the final version of the manuscript.

## ACKNOWLEDGMENT

This work was performed at the University of Rochester and was funded by a grant from the National Institutes of Health, National Institute of Allergy and Infectious Diseases, via grant R61AI167035, and by the Center for Innovation in Point of Care Technologies for HIV/AIDS at Northwestern (sub-award #60058448). Jeffrey W. Beard was supported by an NSF Graduate Research Fellowship. We also thank Prof. David Mathews for helpful comments during the preparation of this manuscript.

## TOC Image

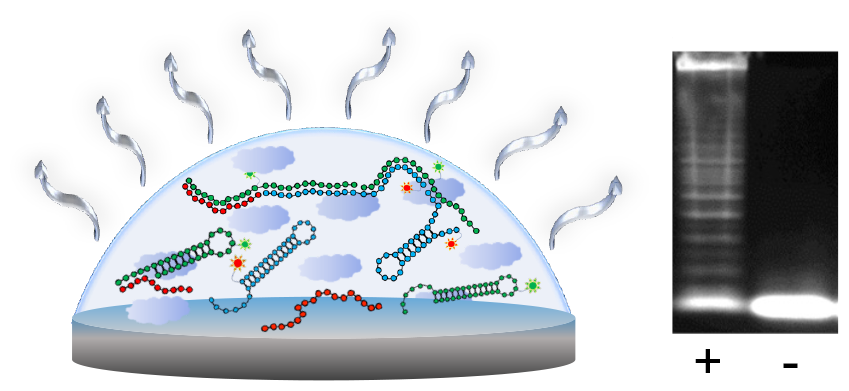

