## Supplementary material for "Mimicking an *in cellulo* environment for enzyme-free paper-based nucleic acid tests at the point of care": Combined Supplementary Information

#### Table of Contents

**Sequences and Tm Experiments………..……………………………….2**

**Functionalized Fusion 5 Experiments……………………………...3**

**Limit of Detection Calculations…………………………………………4**

**HCR with Varying Hairpin Concentrations…………………………...5**

**HCR with Salmon Sperm DNA and Yeast tRNA……………………...6**

**Impact of Tween 20 on Fluorescence…………………………………...7**

**References……………………………………………………………………...8**

### Sequences and Tm Experiments

Table S1. Sequences used in this work. Colors represent complementary regions.

| Name | Sequence | pNL4-3 Genome Location |
|---|---|---|
| DNA Trigger | 5' - CTTCAGGTTTGGGGAAGAGACAACAAC - 3' | 2175-2201 |
| RNA Trigger | 5' - CUUCAGGUUUGGGGAAGAGACAACAAC - 3' | |
| H1 | 5' - GGGGAAGAGACAAGGTTTTGTCTCTTCCCCAAACCT - 3' | |
| H2 | 5' - AAACCTTGTCTCTTCCCCAGGTTTGGGGAAGAGACA - 3' | |
| H1-Cy3 | 5' - GGGGAAGAGACAAGGTTT/iCy3/TGTCTCTTCCCCAAACCT - 3' | |
| H2-Cy5 | 5' - /5Cy5/AAACCTTGTCTCTTCCCCAGGTTTGGGGAAGAGACA - 3' | |
| H1-ATTO550 | 5' - GGGGAAGAGACAAGGTTTTGTCTCTTCCCCAAACCT /ATTO550/ - 3' | |
| H2-ATTO647N | 5' - /ATTO647N/AAACCTTGTCTCTTCCCCAGGTTTGGGGAAGAGACA - 3' | |

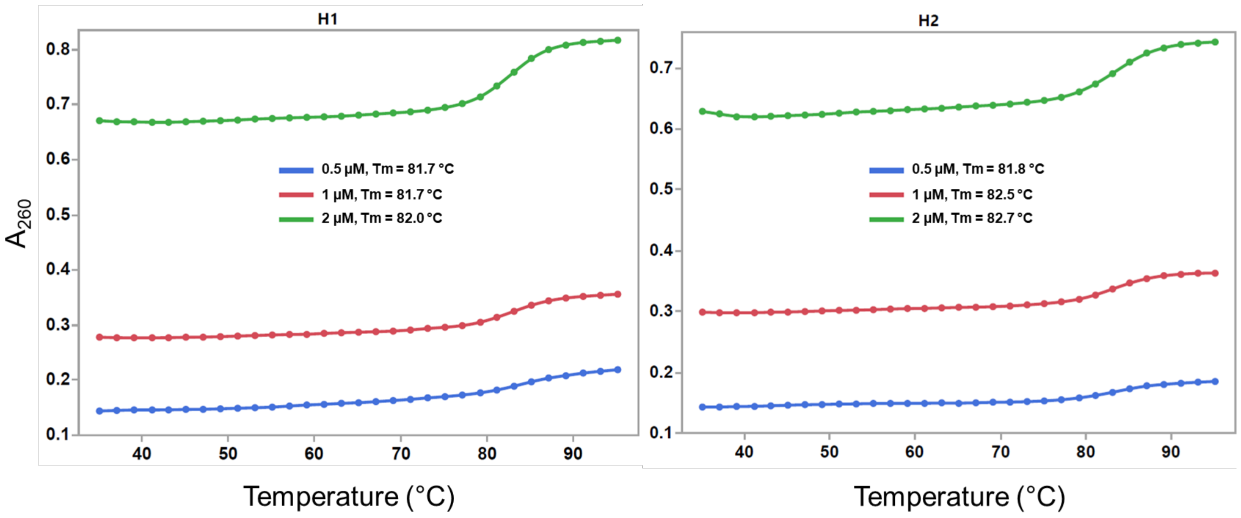


**Figure S1.** Verification of hairpin formation for H1 (left) and H2 (right) hairpins. Protocol described under *Nucleic Acid Oligomers* in Experimental Section.

### Functionalized Fusion 5 Experiments

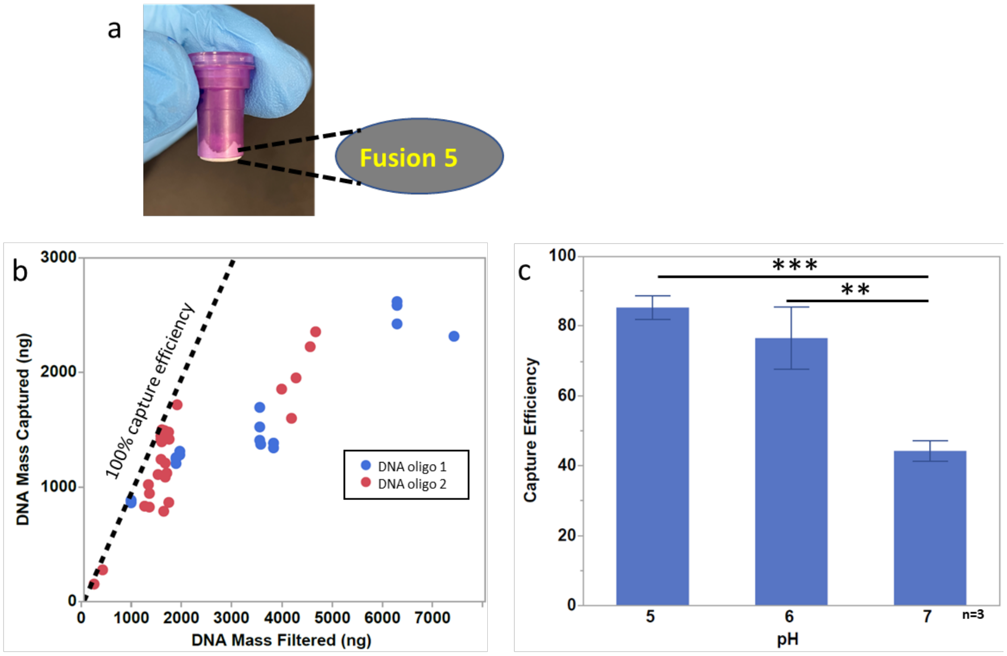


d

| | |
|---|---|
| DNA oligo 1 | CTTCAGGTTTGGGGAAGAGACAAC |
| DNA oligo 2 | AGACTGTAGCCCAGGAATATGGCAGCTAGA |

**Figure S2.** Quantified results of RNA and DNA captured on functionalized Fusion 5 filters. a) Custom centrifuge filter using chitosan-functionalized Fusion 5 filter. b) Mass of DNA oligos 1 and 2 captured on functionalized Fusion 5 compared to the mass of the total DNA filtered. The dashed line represents where 100% capture efficiency of the DNA on the Fusion 5 would occur. c) The capture efficiency of yeast tRNA filtered through the functionalized Fusion 5 in a 1X PBS buffer at pH 5, 6 and 7. d) DNA sequences using in (b).

Based on our experimental data, we estimate the chitosan-functionalized Fusion 5 filter to have a capture capacity of approximately 85 ng/mm$^2$, which is similar to what has been reported by other groups using similar capture filters.[1,2] The 2 mm diameter capture disks used for these experiments can theoretically hold hundreds of nanograms of target nucleic acids. Capture capacity was calculated using data from Figure S2b assuming saturation at 2.4 µg DNA on a 6 mm diameter disc:

$$\frac{2.4\ \mu g}{\pi 3^2} = 85\ \frac{ng}{mm^2}$$

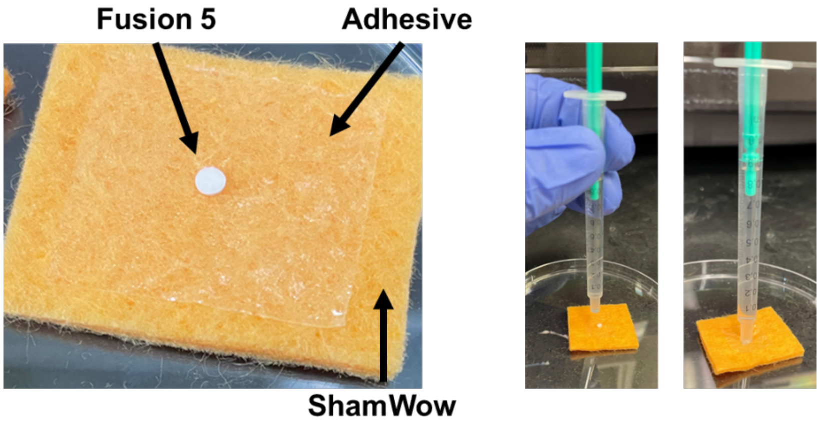


**Figure S3.** Experimental setup for filtering and capturing nucleic acids through chitosan-functionalized Fusion 5. A 2 mm disk of functionalized Fusion 5 is placed in direct contact with a ShamWow absorbent pad by way of double-sided adhesive with a 1.5 mm hole directly under the Fusion 5. A 1 mL syringe filled with the diluted sample was placed firmly in direct contact with the Fusion 5 filter and the syringe plunger was gently depressed (~ 5 seconds to depress plunger fully).

Table S2. Results of DNA filtration shown in Figure S2b

| DNA Used | Centrif. Force (xg) | Time Centrifuged (s) | [DNA](uM) | Volume (uL) | nanograms filtered | nanograms captured | %captured |
|---|---|---|---|---|---|---|---|
| DNA Oligo 1 | 1300 | 60 | 20 | 100 | 1912.7 | 1311.9 | 0.7 |
| DNA Oligo 1 | 1300 | 60 | 20 | 100 | 1912.7 | 1280.7 | 0.7 |
| DNA Oligo 1 | 1300 | 60 | 20 | 100 | 1912.7 | 1280.7 | 0.7 |
| DNA Oligo 1 | 1300 | 60 | 40 | 50 | 1838.1 | 1205.1 | 0.7 |
| DNA Oligo 1 | 1300 | 60 | 40 | 50 | 1838.1 | 1256.6 | 0.7 |
| DNA Oligo 1 | 1300 | 60 | 40 | 50 | 1838.1 | 1245.5 | 0.7 |
| DNA Oligo 1 | 1300 | 60 | 40 | 100 | 3526.1 | 1373.1 | 0.4 |
| DNA Oligo 1 | 1300 | 60 | 40 | 100 | 3781.5 | 1383.3 | 0.4 |
| DNA Oligo 1 | 1300 | 60 | 40 | 100 | 3781.5 | 1341.6 | 0.4 |
| DNA Oligo 1 | 1300 | 60 | 40 | 150 | 6243.5 | 2424.2 | 0.4 |
| DNA Oligo 1 | 1300 | 60 | 40 | 150 | 6243.5 | 2619.0 | 0.4 |
| DNA Oligo 1 | 1300 | 60 | 40 | 150 | 6243.5 | 2586.0 | 0.4 |
| DNA Oligo 1 | 1300 | 60 | 40 | 25 | 942.2 | 861.9 | 0.9 |
| DNA Oligo 1 | 1300 | 60 | 40 | 25 | 942.2 | 879.2 | 0.9 |
| DNA Oligo 1 | 1300 | 60 | 40 | 25 | 942.2 | 884.9 | 0.9 |
| DNA Oligo 1 | 1300 | 60 | 40 | 200 | 7378.8 | 2316.0 | 0.3 |
| DNA Oligo 1 | 1300 | 60 | 40 | 100 | 3506.7 | 1695.1 | 0.5 |
| DNA Oligo 1 | 1300 | 60 | 40 | 100 | 3506.7 | 1524.3 | 0.4 |
| DNA Oligo 1 | 1300 | 60 | 40 | 100 | 3506.7 | 1407.7 | 0.4 |
| DNA Oligo 2 | 4000 | 60 | 10 | 50 | 4620.7 | 2355.4 | 51.0 |
| DNA Oligo 2 | 4000 | 60 | 10 | 50 | 3943.2 | 1854.9 | 47.0 |
| DNA Oligo 2 | 4000 | 60 | 10 | 50 | 4229.7 | 1952.4 | 46.2 |
| DNA Oligo 2 | 4000 | 60 | 10 | 50 | 4512.2 | 2224.8 | 49.3 |
| DNA Oligo 2 | 4000 | 60 | 10 | 50 | 4139.8 | 1599.7 | 38.6 |
| DNA Oligo 2 | 1300 | 60 | 0.25 | 800 | 1212.7 | 832.7 | 68.7 |
| DNA Oligo 2 | 1300 | 60 | 0.5 | 400 | 1305.6 | 824.6 | 63.2 |
| DNA Oligo 2 | 1300 | 60 | 1 | 200 | 1307.7 | 943.9 | 72.2 |
| DNA Oligo 2 | 1300 | 60 | 5 | 40 | 1538.9 | 1241.4 | 80.7 |
| DNA Oligo 2 | 1300 | 60 | 10 | 20 | 1610.4 | 1492.4 | 92.7 |
| DNA Oligo 2 | 1300 | 60 | 25 | 8 | 1654.3 | 1121.2 | 67.8 |
| DNA Oligo 2 | 1300 | 60 | 0.25 | 800 | 1689.7 | 1479.5 | 87.6 |
| DNA Oligo 2 | 1300 | 60 | 0.5 | 400 | 1859.4 | 1718.0 | 92.4 |
| DNA Oligo 2 | 1300 | 60 | 1 | 200 | 1592.6 | 788.2 | 49.5 |
| DNA Oligo 2 | 1300 | 60 | 5 | 40 | 1700.6 | 1417.2 | 83.3 |
| DNA Oligo 2 | 1300 | 60 | 10 | 20 | 1694.1 | 865.4 | 51.1 |
| DNA Oligo 2 | 1300 | 60 | 25 | 8 | 1556.8 | 1499.9 | 96.3 |
| DNA Oligo 2 | 1300 | 60 | 0.5 | 400 | 1552.2 | 1394.6 | 89.8 |
| DNA Oligo 2 | 1300 | 60 | 1 | 200 | 1287.5 | 1020.7 | 79.3 |
| DNA Oligo 2 | 1300 | 60 | 5 | 40 | 1479.9 | 1108.8 | 74.9 |
| DNA Oligo 2 | 1300 | 60 | 10 | 20 | 1536.3 | 1411.6 | 91.9 |
| DNA Oligo 2 | 1300 | 60 | 10 | 20 | 1536.3 | 1444.7 | 94.0 |
| DNA Oligo 2 | 1300 | 60 | 1 | 200 | 1623.0 | 1086.4 | 66.9 |
| DNA Oligo 2 | 1300 | 60 | 1 | 200 | 1623.0 | 1208.6 | 74.5 |
| DNA Oligo 2 | 4000 | 60 | 1 | 50 | 374.8 | 276.3 | 73.7 |
| DNA Oligo 2 | 4000 | 60 | 0.5 | 50 | 205.0 | 152.3 | 74.3 |

### LOD Calculations

**Table S3. Sessile droplet LOD calculations.**

| | **DNA Sessile** | **DNA Tube** | **RNA Sessile** | **RNA Tube** |
|---|---|---|---|---|
| **Experiment 1** | 0.223 | 0.200 | 0.209 | 0.195 |
| **Experiment 2** | 0.215 | 0.199 | 0.211 | 0.180 |
| **Experiment 3** | 0.219 | 0.184 | 0.214 | 0.199 |
| **Standard Deviation (SD)** | 0.004 | 0.009 | 0.003 | 0.010 |

| | | | | |
|---|---|---|---|---|
| **slope (m)** | 0.630 | 0.530 | 0.401 | 0.124 |
| **LOD (nM) = 3.3*SD/m** | **0.019** | **0.057** | **0.021** | **0.271** |

### HCR with Varying Hairpin Concentrations

When incubated with different synthetic DNA trigger concentrations, the normalized FRET intensity of the HCR product from H1-donor and H2-acceptor hairpins increased linearly with trigger concentration up to a 1:1 hairpin:trigger ratio (Figures 2b and 2c). Interestingly, at very high concentrations of trigger (greater than 1:1 hairpin to trigger ratio) this concept breaks down, and we see a dramatic decrease in FRET transmission that asymptotes at a lower intensity than the maximum (Figure S4) This phenomenon results, at least in part, from the placement of the fluorophores (Figure 1a). The first H1 donor and H2 acceptor fluorophores are 36 nucleotides apart due to the placement of the fluorophores on the hairpins resulting in low FRET efficiency. Every subsequent set of fluorophores in the polymer are directly adjacent and approach 100% efficiency. Low concentrations of trigger result in fewer first round H1-H2 hybridization (low FRET efficiency), and more subsequent hybridizations (high FRET efficiency) due to the formation of longer polymer chains. This work prioritizes lower trigger concentration where the effect is negligible.

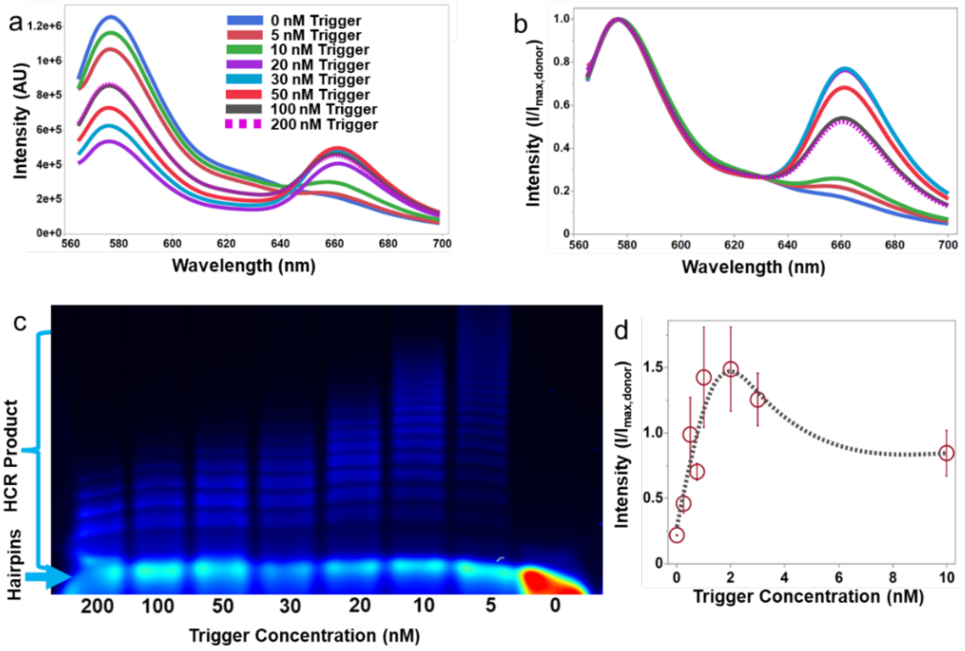


**Figure S5.** Synthetic DNA trigger incubated in cuvettes with 100 nM H1-ATTO550 and H2-ATTO647N hairpins for 1 hour in 5X SSC with 0.05%Tween 20. Panels (a) and (b) show the raw and normalized spectra of each trigger concentration from 0 – 200 nM. The 100 and 200 nM trigger concentrations show nearly identical spectra, indicating that the cascade reaches a saturation point composed of lower molecular weight product (shown in the Cy5 channel of the agarose gel in panel (c)) at a 1:1 ratio of hairpin to trigger. d) 5 nM H1-Cy3 and H2-Cy5 hairpin concentration incubated with synthetic DNA trigger in sessile droplets for 1 hour in 0.5X SSC with 0.1% Tween 20 and 10 mM $MgCl_2$. There is a decrease in intensity that levels off around 5 nM trigger concentration, consistent with the 1:1 hairpin to trigger ratio described at other hairpin concentrations. Statistical data was analyzed using JMP Pro 16. Error bars show standard deviation from an n = 3.

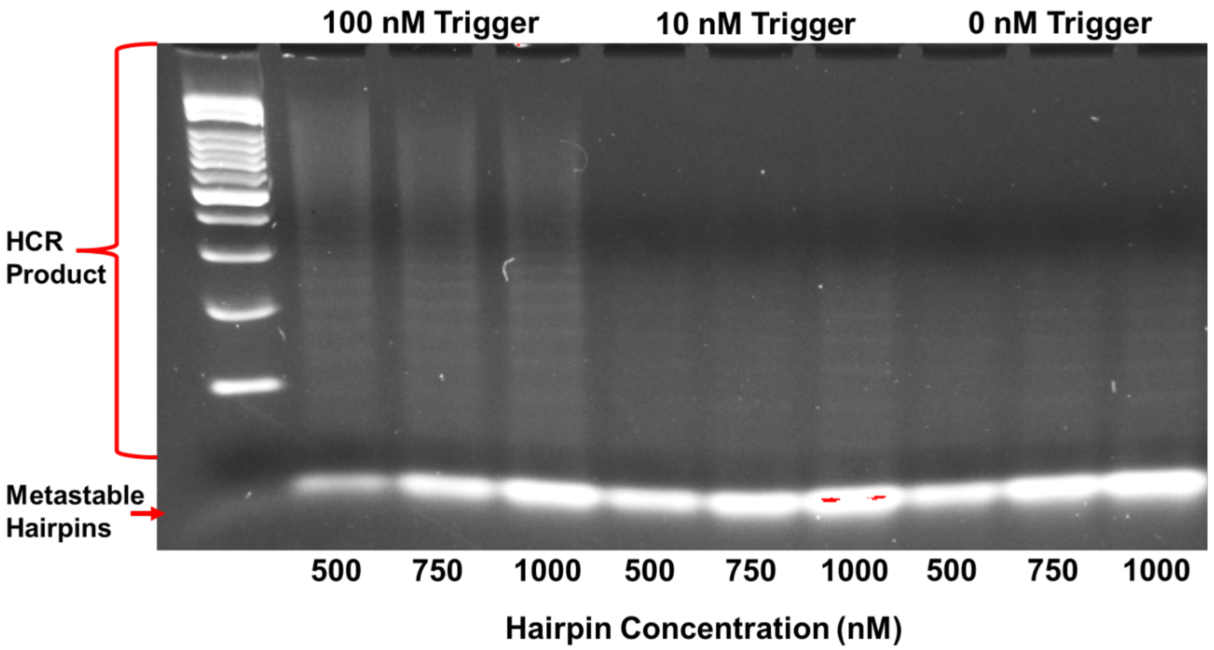


**Figure S6**. Increasing hairpin concentrations with fixed synthetic RNA trigger concentrations. Hairpins at 500, 750, and 1000 nM concentrations were incubated in 5X SSC with 0.05% Tween 20 with 100, 10, and 0 nM synthetic RNA trigger concentrations. The first lane is a 100 base pair molecular weight ladder.

### HCR with Salmon Sperm DNA and Yeast tRNA

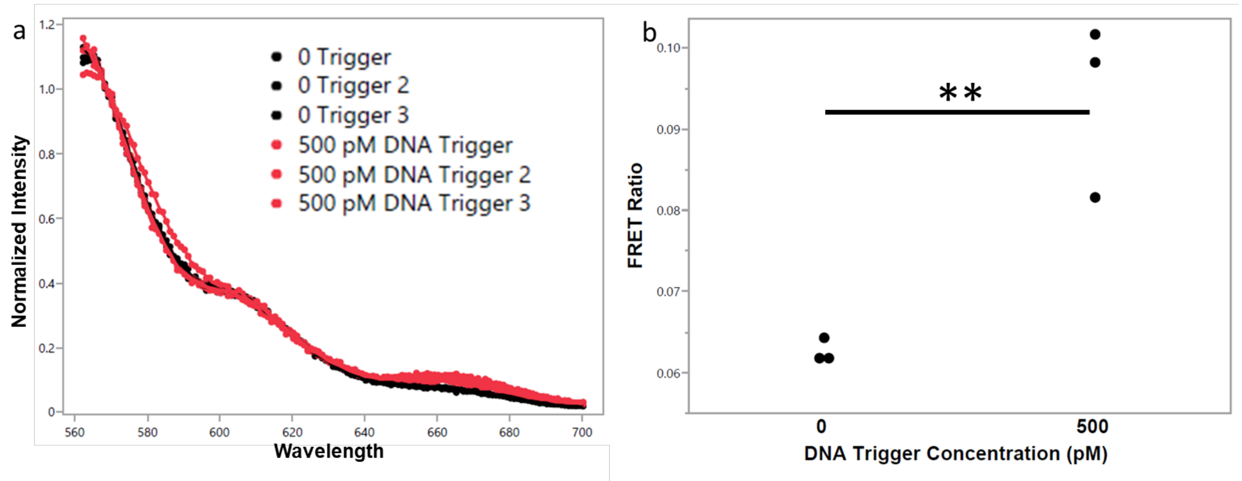


**Figure S7.** H1-Cy3 and H2-Cy5 incubated with salmon sperm DNA in sessile droplets for 1 hour in 0.5X SSC with 0.1% Tween 20 and 10 mM $MgCl_2$. a) Normalized spectra of the results with (red) and without (black) the presence of DNA trigger, and b) the FRET ratio of the results. Each 5 µL reaction contained 25 ng of salmon sperm DNA. The reactions with trigger contained approximately 22 pg of DNA trigger per reaction, for approximately 1:1000 ratio of trigger to salmon sperm DNA. This data was collected using a Fluoromax Plus (Horiba, Kyoto, Japan), a different instrument from the one used to collect the spectral data shown in the main text. Statistical data was analyzed using JMP Pro 16. Error bars show standard deviation from an n = 3, and ** $p < 0.01$.

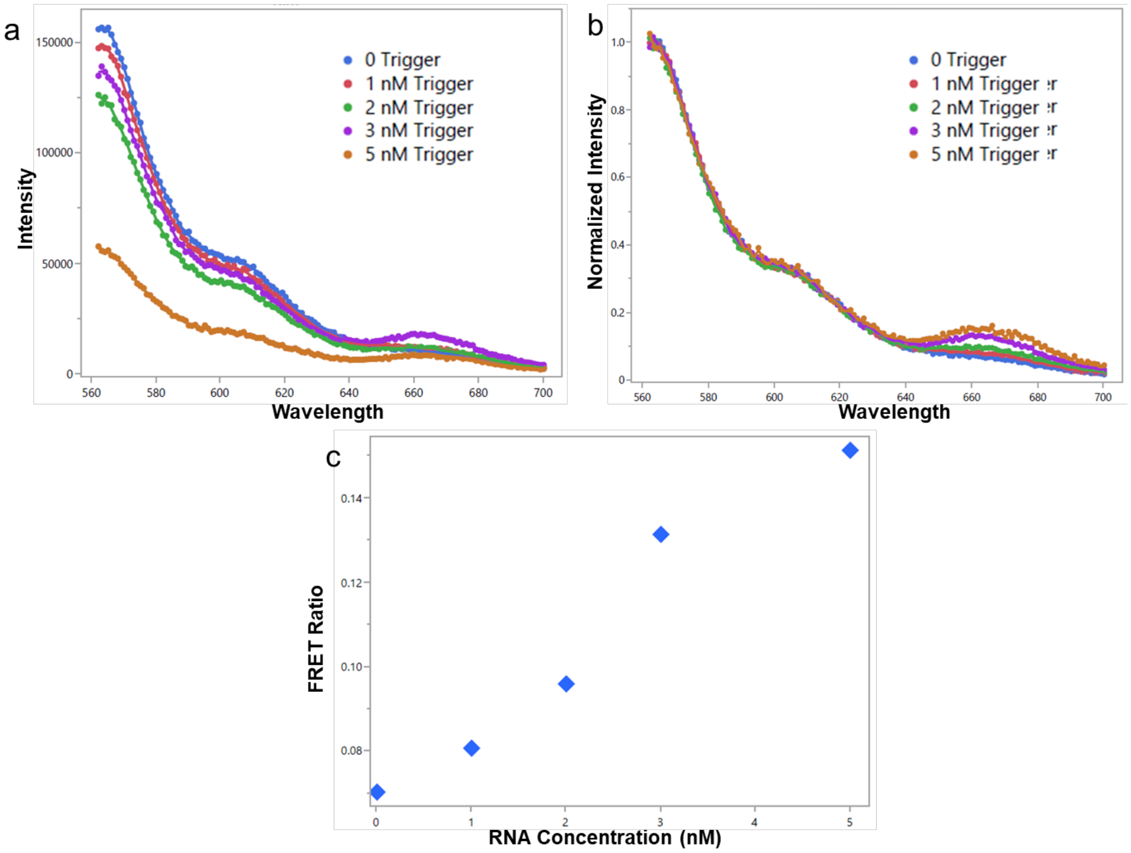


**Figure S8**. 1 µg/mL of Yeast tRNA incubated with H1-Cy3 and H2-Cy5 in sessile droplets for 1 hour in 0.5X SSC with 0.05% Tween 20 and 10 mM $MgCl_2$. a) and b) The spectra and normalized spectra of the results when incubated with varying concentrations of synthetic RNA trigger. c) Plot of each FRET ratio versus trigger concentration (n=1). This data was collected using a Fluoromax Plus (Horiba, Kyoto, Japan), a different instrument from the one used to collect the spectral data shown in the main text.

### Impact of Tween 20 on Fluorescence

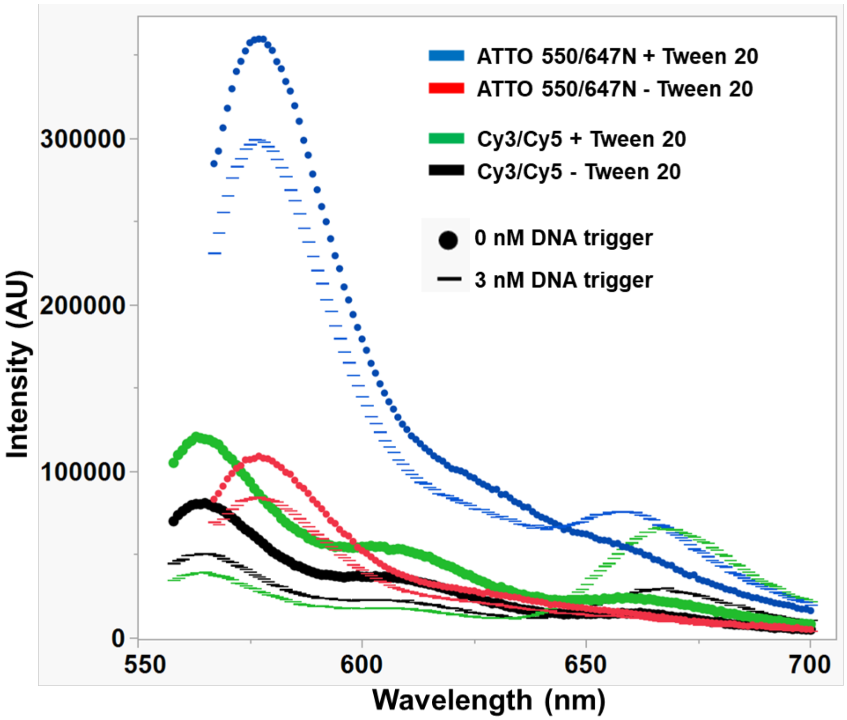


**Figure S9.** Impact of Tween 20 on fluorescence output of Cy- and ATTO dyes. 5 nM concentrations of fluorescently tagged H1 (Cy3 or ATTO 550) and H2 (Cy5 or ATTO 647N) were incubated in 5X SSC buffer with and without the addition of 0.05% Tween 20. The reactions were incubated with and without 3 nM synthetic DNA trigger for 1 hour at room temperature in cuvettes.
